# Viral characteristics associated with sustenance of elite neutralizing activity in chronically HIV-1C infected monozygotic pediatric twins

**DOI:** 10.1101/475822

**Authors:** Nitesh Mishra, Muzamil Ashraf Makhdoomi, Shaifali Sharma, Sanjeev Kumar, Deep-shika Kumar, Himanshi Chawla, Ravinder Singh, Uma Kanga, Bimal Kumar Das, Rakesh Lodha, Sushil K Kabra, Kalpana Luthra.

**Affiliations:** Department of Biochemistry, All India Institute of Medical Sciences, New Delhi, India.; Department of Biochemistry, Government College for Women, Cluster University Srinagar, Srinagar, India.; Department of Microbiology, All India Institute of Medical Sciences, New Delhi, India.; Department of Transplant Immunology and Immunogenetics, All India Institute of Medical Sciences, New Delhi, India.; Department of Pediatrics, All India Institute of Medical Sciences, New Delhi, India.

**Keywords:** Epitope mapping, HIV-1C envelope, pediatric HIV-1C infection, broadly neutralizing antibodies, V2-glycan, V3-glycan, pediatric elite neutralizers.

## Abstract

Broad and potent neutralizing antibodies (bnAbs) with multiple epitope specificities evolve in HIV-1 infected children. Herein we studied two antiretroviral naïve chronically HIV-1C infected monozygotic pediatric twins AIIMS_329 and AIIMS_330 with potent plasma bnAbs. Elite plasma neutralizing activity was observed since initial sampling at 78 months in AIIMS_330 and persisted throughout, while in AIIMS_329 it was seen at 90-months of age after which potency decreased overtime. We evaluated potential viral characteristics associated with the varied immune profile by generating single genome amplified pseudoviruses. The AIIMS_329 viruses generated from 90-month time point sampling were neutralization sensitive to second generation bnAbs and contemporaneous autologous plasma antibodies, while viruses from 112-months and 117-month timepoints were resistant to most bnAbs and autologous contemporaneous plasma. AIIMS_329 viruses developed resistance to plasma nAbs plausibly by N160-glycan loss, V1- and V4-loop lengthening. The viruses generated from AIIMS_330 at 90-month and 117-month timepoint showed varied susceptibility to bnAbs and autologous contemporaneous plasma antibodies while the viruses of 112-month timepoint, at which plasma nAb specificities mapped to the V2-glycan, V3-glycan and CD4bs, were resistant to autologous contemporaneous plasma antibodies as well as most bnAbs. We observed evolution of a viral pool in AIIMS_330 donor, comprising of plasma antibody neutralization sensitive or resistant diverse autologous viruses that in turn may have contributed to development and maintenance of elite neutralizing activity. The findings of this study provide information towards understanding factors involved in generation and maintenance of potent plasma nAbs.

**Importance:** Chronically HIV-1 infected children that develop elite neutralizing activity are suitable candidates to understand the mechanisms that lead to the co-evolution of virus and antibody response. Here, we evaluated the alterations in virus and antibody responses over time in chronically HIV-1C infected monozygotic pediatric twins, AIIMS_329 and AIIMS_330, who had acquired the infection by vertical transmission. AIIMS_330 retained the elite plasma neutralizing activity throughout, while in AIIMS_329, the potency decreased post 90 months of age. The corresponding viral pool from post 90-month samples in AIIMS_330 showed varied susceptibility, while that in AIIMS_329, developed resistance to bnAbs and autologous plasma antibodies. The findings of this study, conducted in twin children of same genetic make-up and infected at birth with a single source of HIV-1C, suggest that a viral pool with varied susceptibility to antibodies could have been one of the factors responsible for sustained elite neutralizing activity in AIIMS_330.

## Introduction

Disease progression in HIV-1 infected children is observed to develop in a biphasic manner. More than 50% of the infected children, if not initiated on antiretroviral therapy (ART), progress to AIDS within a couple of years of infection due to an immature immune system and higher levels of viral replication (1,2). Infected children who survive beyond the initial years mostly develop chronic disease (3) with slow progression. HIV-1 infected children have been shown to mount potent plasma broadly neutralizing antibodies (bnAbs) earlier than adults and more frequently with multiple bnAb specificities. than adults (4–6). The bnAb BF520.1 was recently isolated from an infant one-year post-infection (p.i) (4,7). The presence of epitopes, targeted by the second generation bnAbs, on the functional native like HIV-1 envelope trimers identifies such envelopes as candidates for HIV-1 immunogen design. The native-like HIV-1 gp140 trimeric envelope glycoprotein BG505.T332.SOSIP.664.C2 T332N (8–10), generated from the circulating virus of a clade A infected infant, is the best immunogen documented so far. A recent study on 303 HIV-1 transmission pairs showed that among majority of infecting viral strains, only few had the ability to induce bnAb responses (11). Tracing the virological characteristics of such rare viral antigens capable of initiating and shaping potent antibody responses to HIV-1 will provide key insights for future vaccine strategies capable of eliciting similar bnAb responses (11,12). Thus, pediatric HIV-1 infection, with its unique characteristics of rapid and frequent induction of potent bnAbs with multiple specificities, provides a unique model to understand the virological characteristics that can be utilized for future HIV-1 vaccine candidates capable of inducing bnAbs.

Chronically infected children have been shown to have potent plasma bnAbs with diverse epitope specificities (5,6). In our pediatric cohort of antiretroviral naïve HIV-1C infected long term non-progressors (LTNPs) characterized for plasma antibody responses (6,13–18), we identified a pair of antiretroviral naive HIV-1C infected children AIIMS_329 and AIIMS_330, defined herein as genetically identical twins by their identical HLA haplotypes, who had acquired the infection at birth by vertical transmission. Both twins AIIMS_329 and AIIMS_330 developed potent plasma neutralizing antibodies, with the latter showing elite neutralizing activity since first sampling. Elite neutralizers are a rare subset of LTNPs that constitute the top 1% of HIV-1 infected individuals with ability to neutralize pseudoviruses belonging to multiple clades at ID50 titres of 300 or more. A systematic analysis of the circu-lating viral variants in elite neutralizers over time, their antibody imprinting ability (11) and correlation with evolving autologous plasma neutralization response (19), can provide useful information for immunogen design. A study conducted on HIV-1 infected identical and fraternal twins to evaluate for variation in the C2-C5 viral sequence or viremia showed that in majority of the twin pairs, minimal sequence variation was seen in the early viruses indicating that they were infected by the same or closely related quasispecies from the mother. Over time, the viruses showed divergence. The similar viral levels observed in these twins further suggested that inherent viral properties determine the pattern of viral expression with a minimal role of host genetics (20).

We evaluated AIIMS_329 and AIIMS_330 for a period of 60 months, with time matched sampling since their baseline sampling in 2013, to decipher the viral characteristics responsible for the induction and sustenance of potent plasma bnAbs. One of the twins, AIIMS_330, showed elite neutralizing activity at first sampling that improved with time. The AIIMS_330 plasma showed polyclonal bnAb response with multiple bnAb specificities that increased in potency in the follow up time points, and had multiple circulating viral variants with varied susceptibility to contemporaneous autologous plasma antibodies and bnAbs. The plasma of the second twin AIIMS_329, also developed elite neutralizing activity, although at a later time point. In order to determine the virological traits associated with the diverse plasma bnAb response, we generated envelope pseudoviral clones of the circulating viruses from the twin plasma samples at different time points and tested their susceptibility to neutralization by the contemporaneous and evolving plasma antibodies and to the known bnAbs to identify the neutralizing determinants on these viruses of pediatric origin. AIIMS_329 showed the presence of circulating viral variants that were resistant to autologous contemporaneous plasma and bnAb neutralization. Taken together, our data shows the presence of multiple and distinct circulating viral variants, with varied susceptibility to autologous plasma antibodies and bnAbs in each of the two genetically identical HIV-1 infected twins, that impacted the varied evolution of polyclonal bnAb response despite the similar genetic background in the twins.

## Results

### Varied evolution of plasma antibodies in genetically identical twins against diverse clades of HIV

In this study, we evaluated a total of nine independent and time-matched plasma samples from AIIMS_329 and AIIMS_330 pediatric HIV-1C infected twin donors, earlier established as LTNPs (6). High resolution HLA typing revealed that AIIMS_329 and AIIMS_330 were identical at HLA-A, B, DRB1 loci, and contained wild type CCR5 alleles (data not shown). The detailed two field HLAs were HLA-A*02:11, *24:02; B*35:03*40:06; DRB1*14:04*15:01. The baseline (First) sampling of the identical twins was done at the age of 78-months, and they were then longitudinally followed up for a period of 60 months upto the age of 138-months. The CD4+ T cell counts and viral loads of respective time-points are given in Table 1.

**Table 1:**
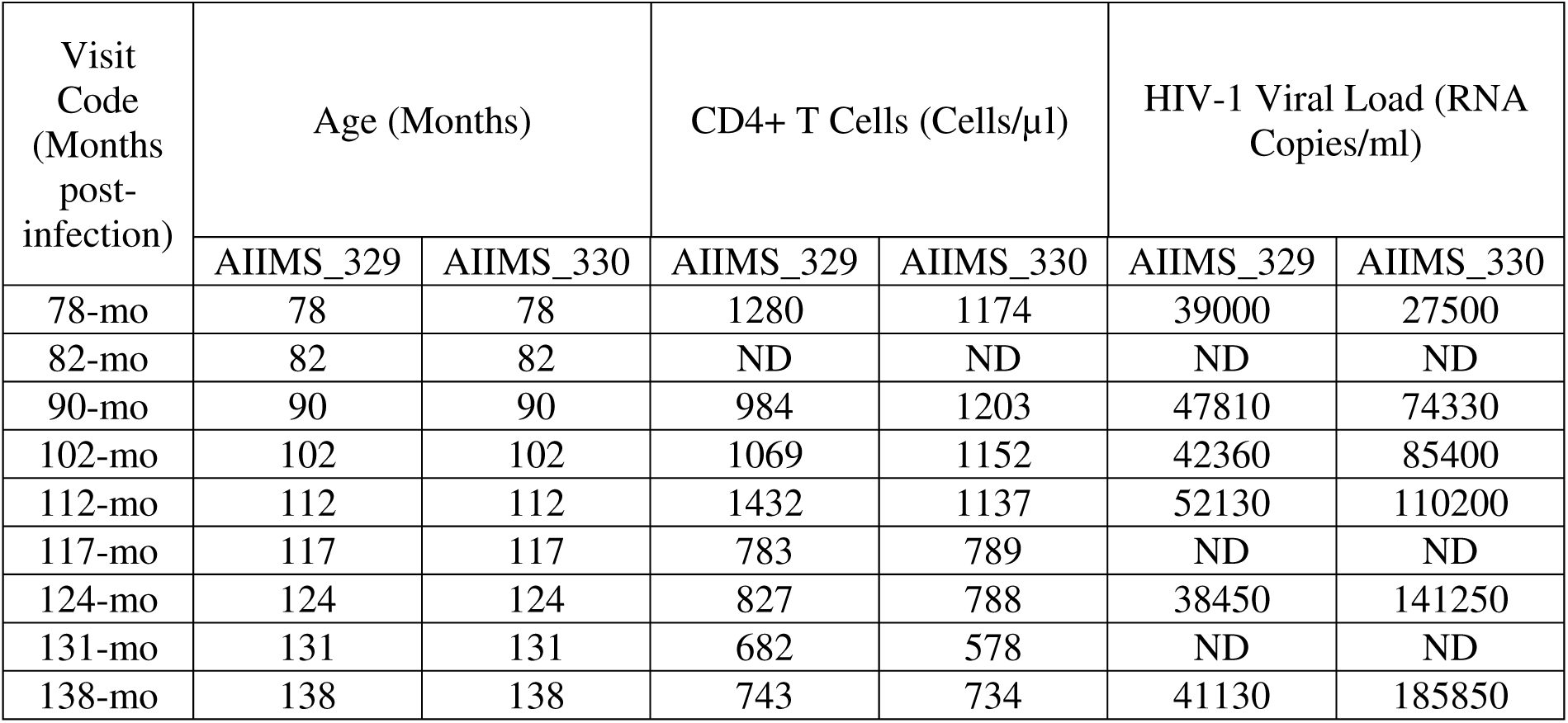
Demographic and clinical characteristics of AIIMS_329 and AIIMS_330. ND – Not Determined.

A longitudinal assessment of evolution of plasma antibodies of AIIMS_329 and AIIMS_330 was done over a period of 60 months utilizing a standardized panel of 12 HIV-1 envelope pseudoviruses that represent strains circulating across the globe (21). The HIV-1 neutralizing activity of plasma antibodies was assessed by performing neutralization assays using TZM-bl based luciferase reporter assay. At the 78-month sample, the plasma antibodies of both the twins showed potent neutralizing activity against most of the pseudoviruses in the panel (10 out of 12 for AIIMS_329 and 9 out of 12 for AIIMS_330) (**Figure 1A and C**). The AIIMS_330 plasma antibodies demonstrated increase in neutralization breadth (from 75% to 100% viruses neutralized) and potency (GMT titres from 204 to 929) over time, and reached a maximum ID50 titre of 966 at 117-months, that persisted till the last sample time point (**Figure 1C-D**). Of particular note was the neutralization of majority of viruses at plasma dilution of 300 or more. In case of AIIMS_329, the potency (GMT titres from 244 to 384) increased up to 90 months, followed by a reduction in neutralizing antibody potency (GMT titres of 93), that stabilized at 112 months till the final sample time point (GMT titre 124). (**Figure 1A-B**). There was a reduction in AIIMS 329 plasma antibody ID50 titre against non-clade C viruses, whereas the titre against 25710_2_43 pseudovirus (Indian clade C), despite showing a reduction, was maintained in the range of ID50 300-700.

**Figure 1:**
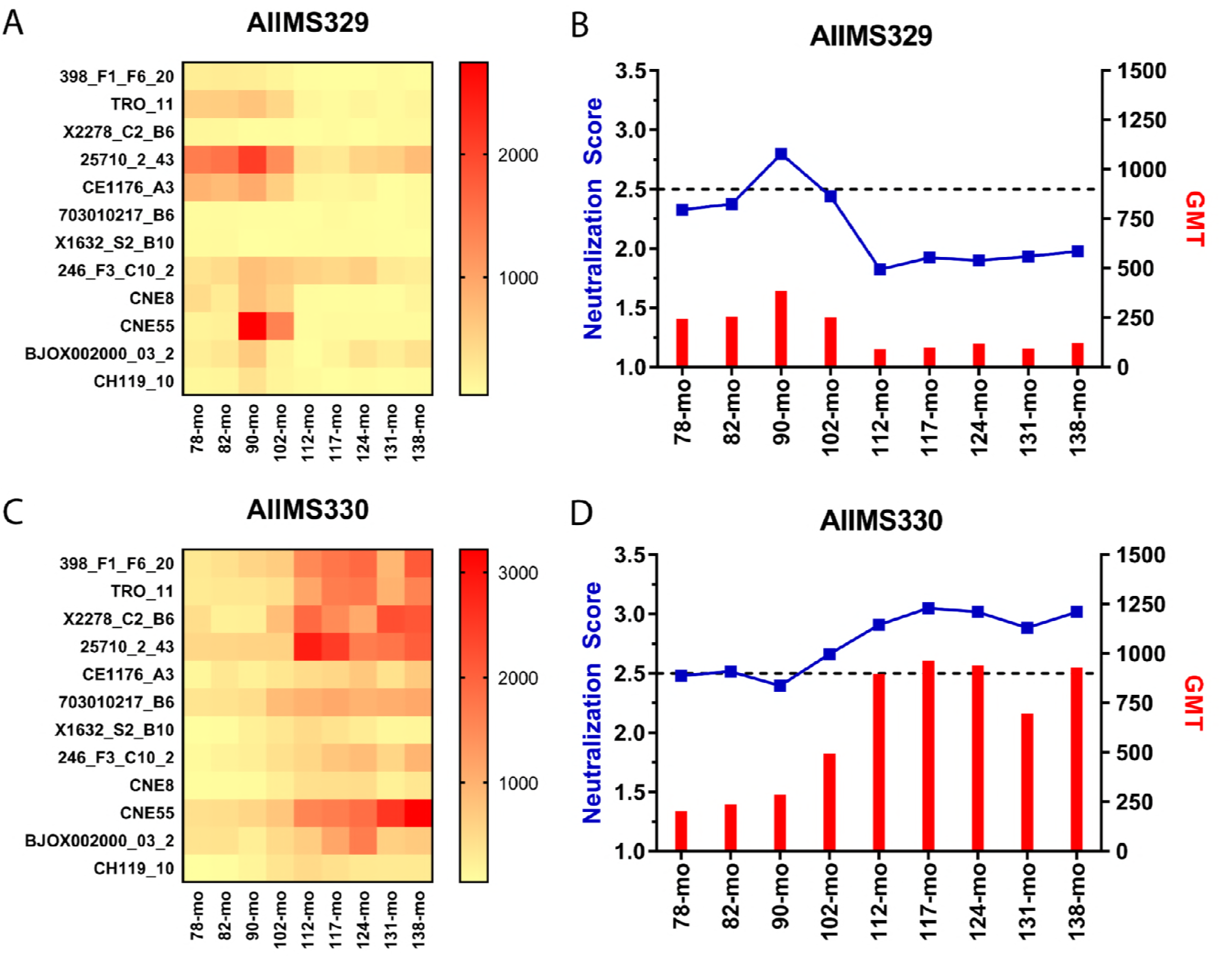
Plasma neutralization breadth of AIIMS_329 and AIIMS_330 against the 12-virus global panel. (A and C) Heatmap showing the longitudinal plasma ID50 values, shown as reciprocal plasma dilution from 3 independent experiments, of AIIMS_329 and AIIMS_330. The horizontal bar shows the color key. (B and D) longitudinal evolution of GMT titres and neutralization scores of plasmas of AIIMS_329 and AIIMS_330. A cut-off of 2.5 signifies elite neutralizing activity.

To substantiate this observation and to evaluate for elite neutralizing activity, we further assessed plasma neutralizing activity using an extended multiclade 45 virus panel. With the extended panel of viruses, viral neutralization was checked with plasma follow up time points until 112 months, since both the twins showed ability of plasma antibodies to neutralize the global panel of viruses beyond the age of 112 months. AIIMS_329 plasma showed elite neutralizing activity at 90-months after which the plasma potency decreased overtime, against non-clade C viruses (**Figure 2A-B**). As observed with the 25710 clade C pseudovirus in the global panel, AIIMS_329 plasma maintained its neutralization activity against the tested clade C viruses of Indian origin, unlike the reduction seen in the ability to neutralize non-clade C viruses (**Figure 2B**). AIIMS_330 plasma in its baseline sampling at 78-months showed elite neutralizing activity against the 45-virus panel, i.e., with ability to neutralize more than one pseudovirus at a dilution of 300 or more within a clade group and across at least four clade groups (**Figure 2C**). For AIIMS_330, unlike AIIMS_329, the elite neutralizing activity consistently increased in potency overtime till the final time point sampling, with the most noteworthy increase in titres observed at 112 months. As we had observed a plateau in ID50 titres beyond this time point with the global panel of viruses, we did not perform further neutralization assays of samples after 112 months of follow up.

**Figure 2:**
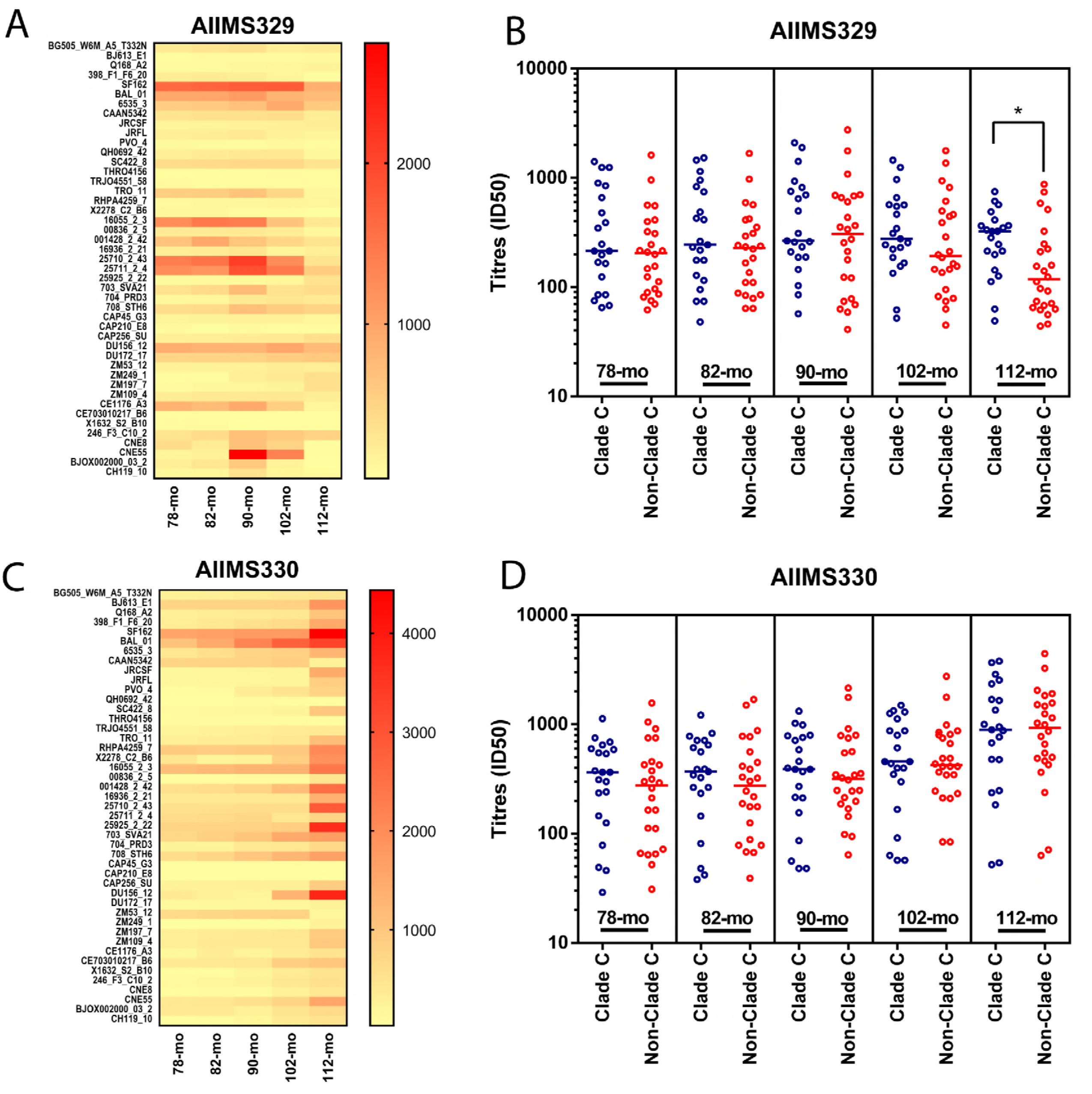
Plasma neutralization breadth of AIIMS_329 and AIIMS_330 against the multiclade 45-virus panel. (A and C) Heatmap of longitudinal ID50 values of AIIMS_329 and AIIMS_330 against the extended 45-virus panel. (B and D) Comparison of clade C and non-clade C ID50 titres of AIIMS_329 and AIIMS_330 from 78-months to 112-months. Mann-Whitney U test was used to compare the ID50 titres of clade C vs non-clade C viruses.

### AIIMS_330 developed antibodies targeting V2-glycan, V3-glycan and CD4bs epitopes with increase in potency of V2-glycan antibodies with time

In order to identify the neutralizing specificities of AIIMS_329 and AIIMS_330 plasma anti-bodies that may have contributed to their potent neutralizing activity, we tested plasma neutralization of mutant pseudoviruses harbouring mutations at key residues of V2-glycan and V3-glycan to assess glycan dependence (5,6), binding reactivity with RSC3 wild type probe and its mutant RSC3Δ371I/P363N (22) for presence of CD4bs dependence and MPER peptides (MPER-B and –C) (23,24) for MPER dependence.

To test for V2 glycan dependence, we used N160A mutants of BG505 and 16055 pseudo-viruses. Dependence was assigned when neutralizing activity for both the mutant viruses were at least 3-fold lesser than that of the corresponding wild type virus. Based on these criteria, V2 glycan directed neutralizing antibodies were found in AIIMS_330 plasma at all time points including the baseline, while AIIMS_329 showed the emergence of V2 glycan directed plasma neutralizing antibodies at 102-months, the timepoint at which AIIMS_329 plasma antibodies demonstrated elite neutralizing activity (Table 2). In both twins, plasma V2 glycan dependent neutralizing activity increased with time. As V2-glycan antibodies also target residues other than N160 in the V1 and V2 loops, we next assessed the dependence on N156, K169, and K171 residues using alanine mutants of 16055 virus, and both the twins did not show any dependence on N156, K169, and K171, suggesting that plasma V2-glycan antibodies depended solely on N160 (Table 2). Testing for the N332 dependence using the N332A mutants of CAP256 and BG505 N332 viruses showed that V3 supersite glycan directed nAbs evolved 117-months onwards in the plasma of both twins (Table 2).

**Table 2:** V2- and V3-glycan specificity of AIIMS_329 and AIIMS_330 plasma. Both the twins developed neutralizing antibodies against V2-glycan and V3-glycan. AIIMS_329 developed V2-glycan antibodies at 102-months while AIIMS_330 showed V2-glycan antibodies from the baseline sample of 78-months. Both the twins developed V3-glycan antibodies at 117-months.

|  | **Timepoint** | **BG505**<br>**N160A** | **16055**<br>**N160A** | **16055**<br>**N156A** | **16055**<br>**K169A** | **16055**<br>**K171A** | **CAP256**<br>**N332A** | **BG505**<br>**N332T** |
| --- | --- | --- | --- | --- | --- | --- | --- | --- |
| AIIMS_329 | 78-mo | 1.3 | 2.7 | 1.1 | 1 | 1.1 | 1.2 | 0.8 |
|  | 82-mo | 1.1 | 2.9 | 1.1 | 1.1 | 1.4 | 0.9 | 1 |
|  | 90-mo | 1.1 | 2.8 | 1 | 1.1 | 1.2 | 1.1 | 1 |
|  | 102-mo | **3.1** | **4.1** | 1.2 | 0.9 | 1.1 | 1.1 | 1.2 |
|  | 112-mo | **3.4** | **4.3** | 1.3 | 1.1 | 0.9 | 1.2 | 1.2 |
|  | 117-mo | **14** | **8.9** | 1.1 | 1 | 0.9 | **8.3** | **3.2** |
|  | 124-mo | **5.8** | **5.9** | 1 | 1.2 | 1.1 | **5.3** | **3.9** |
|  | 131-mo | **7.8** | **6.8** | 1 | 1.4 | 1.2 | **4.2** | **5.6** |
|  | 138-mo | **9.8** | **8.9** | 1.1 | 1.1 | 1 | **6.1** | **7.4** |
|  | **Timepoint** | **BG505**<br>**N160A** | **16055**<br>**N160A** | **16055**<br>**N156A** | **16055**<br>**K169A** | **16055**<br>**K171A** | **CAP256**<br>**N332A** | **BG505**<br>**N332T** |
| AIIMS_330 | 78-mo | **4.7** | **5** | 1.1 | 1 | 1.1 | 1.1 | 1 |
|  | 82-mo | **4.3** | **5.3** | 1.2 | 1 | 1.2 | 1.3 | 1.1 |
|  | 90-mo | **4.1** | **4.4** | 1.2 | 1.2 | 1.1 | 1.4 | 1.3 |
|  | 102-mo | **4.2** | **4.5** | 1.3 | 1.1 | 0.9 | 1.6 | 1.5 |
|  | 112-mo | **4.7** | **5.2** | 1.1 | 1.3 | 0.9 | 1.7 | 1.6 |
|  | 117-mo | **19** | **5.7** | 0.9 | 0.9 | 1 | **9.8** | **5.3** |
|  | 124-mo | **15.8** | **3.3** | 0.9 | 1 | 1.4 | **6.4** | **4.1** |
|  | 131-mo | **17** | **9.6** | 0.8 | 1 | 1.5 | **5.6** | **6.4** |
|  | 138-mo | **15.4** | **13.4** | 1.1 | 1.1 | 1.4 | **7.4** | **7.8** |

The presence of CD4bs directed antibodies was determined by performing ELISAs using wild type RSC3 protein and its mutant RSC3Δ371I/P363N. The CD4bs directed bnAb VRC01 was used as a positive control. The plasma of a healthy HIV-1 seronegative donor and 447-52D mAb were used as negative controls. A reduction of three-fold or more in binding to RSC3Δ371I/P363N compared to RSC3 was scored as positive for the presence of CD4bs directed antibodies. AIIMS_330 plasma showed presence of CD4bs antibodies at 112- and 117-months, which at later timepoints waned and no CD4bs dependence was seen at 131 and 138 months (Table 3). AIIMS_329 did not show a significant difference in binding to RSC3 and RSC3Δ371I/P363N at all timepoints tested, indicating the absence of CD4bs directed antibodies in the plasma. For presence of antibodies against MPER, ELISA binding assay using MPER peptides was used. The MPER directed bnAb 2F5, was used as a positive control and an HIV-1 seronegative healthy donor plasma was used as negative control. We did not observe MPER directed plasma antibodies in any of the time points tested for both the twin plasma samples (Data not shown).

**Table 3:** Mapping of CD4bs antibodies in the plasma of AIIMS_329 and AIIMS_330. Plasma was serially diluted and tested for binding to RSC3 protein and its mutant RSC3Δ371I/P363N by ELISA. Given are the OD values at 450 nm at plasma dilution of 1:100. VRC01, a CD4bs bnAb isolated using RSC3 probe, was used as positive control.

|  | AIIMS_329 |  |  | AIIMS_330 |  |  | VRC01 |  |  |
| --- | --- | --- | --- | --- | --- | --- | --- | --- | --- |
|  | RSC3 | RSC3Δ371I/P363N | Fold Change | RSC3 | RSC3Δ371I/P363N | Fold Change | RSC3 | RSC3Δ371I/P363N | Fold Change |
| 78-mo | 0.248 | 0.316 | 0.78 | 0.195 | 0.246 | 0.79 | 2.411 | 0.143 | **16.86** |
| 82-mo | 0.298 | 0.246 | 1.21 | 0.326 | 0.294 | 1.11 |  |  |  |
| 90-mo | 0.198 | 0.132 | 1.5 | 0.224 | 0.199 | 1.13 |  |  |  |
| 102-mo | 0.352 | 0.162 | 2.18 | 0.448 | 0.166 | **2.7** |  |  |  |
| 112-mo | 0.322 | 0.205 | 1.57 | 0.579 | 0.139 | **4.16** |  |  |  |
| 117-mo | 0.295 | 0.196 | 1.51 | 0.626 | 0.201 | **3.11** |  |  |  |
| 124-mo | 0.567 | 0.529 | 1.07 | 0.541 | 0.502 | 1.07 |  |  |  |
| 131-mo | 0.571 | 0.461 | 1.24 | 0.474 | 0.403 | 1.18 |  |  |  |
| 138-mo | 0.417 | 0.426 | 0.98 | 0.455 | 0.644 | 0.71 |  |  |  |

### Autologous resistant viruses of both the twins showed distinct mechanisms to escape contemporaneous plasma antibodies as well as bnAbs

As seen in plasma neutralization and mapping assays, the most striking differences observed in both the twins were in the timeframe of 90-months to 117-months with a distinct reduction in AIIMS_329 and increase in AIIMS_330 plasma neutralization potency, and dependence on multiple epitopes. In order to determine the potential viral characteristics associated with the varied immune profile in both the twins, we generated pseudoviruses by single genome amplification (SGA) of the envelope gene from HIV-1 RNA, isolated from the plasma. Further, all the amplicons generated by SGA were directly sequenced to assess the viral diversity. At least 20 amplicons from each timepoint were sequenced to give a 90% confidence interval, to cover most of the circulating strains. The sequences of the viral amplicons were checked for their clade (**REGA HIV Subtyping Tool**), pairwise distance to reference sequence (**Phylo-Place**) and co-receptor usage (**Web PSSM**). All amplicons belonged to Clade C, had the highest phylogenetic relatedness to the reference sequence C.IN.95.95IN21068.AF067155 with a median pairwise sequence distance of 0.093, and utilized CCR5 as the co-receptor. All the AIIMS_329 viruses generated from the 90-month time point sampling were susceptible to neutralization by the majority of the second generation bnAbs as well as contemporaneous autologous plasma antibodies. The pseudoviruses generated from 112-months and 117-month timepoints were resistant to most of the bnAbs and less susceptible to neutralization by the autologous contemporaneous plasma (Table 4). The viruses generated from AIIMS_330 from 90 and 117-months, except for one pseudovirus from 117-mo timepoint showed varied susceptibility to bnAbs and were susceptible to neutralization by autologous contemporaneous plasma antibodies. The AIIMS_330 pseudoviruses of 112-month timepoint, at which plasma neutralizing antibodies specificities mapped to the V2-glycan, V3-glycan and CD4bs, were less susceptible to neutralization by most bnAbs and resistant to autologous contemporaneous plasma antibodies as well as (Table 4).

**Table 4:** Neutralization susceptibilities of envelope pseudoviruses prepared from the plasmas of both the twins to broadly neutralizing antibodies recognizing major epitopes of viral envelope and contemporaneous autologous plasma. The IC50 of bnAbs and ID50 of plasmas were determined in TZM-bl based neutralization assay by titrating the pseudoviruses with serially diluted bnAbs and plasma.

|  |  |  | V2-Glycan |  |  |  | V3-Glycan |  | CD4bs | Interface | MPER | Autologous Plasma |
| --- | --- | --- | --- | --- | --- | --- | --- | --- | --- | --- | --- | --- |
|  |  |  | PG9 | PG16 | PGT145 | CAP256.09 | AIIMS-P01 | PGT121 | VRC01 | PGT151 | 10-E8 |  |
| AIIMS_329 | 90-mo | 329.14.B1 | <0.02 | 5.37 | >10 | >10 | 0.08 | <0.02 | 0.59 | 3 | 3 | 293 |
|  |  | 329.14.B4 | 0.35 | 4.59 | >10 | >10 | 0.17 | 0.05 | 1.25 | 2.98 | 3.74 | 312 |
|  | 112-mo | 329.15.A1 | >10 | >10 | >10 | >10 | 0.48 | >10 | 5.62 | 4.59 | 1.06 | 149 |
|  |  | 329.15.B1 | >10 | >10 | >10 | >10 | 2.94 | 1.26 | >10 | 1.89 | 0.99 | 135 |
|  | 117-mo | 329.16.C1 | >10 | >10 | >10 | >10 | >10 | 2.35 | 2.56 | >10 | 2.11 | <100 |
|  |  | 329.16.D5 | >10 | >10 | >10 | >10 | >10 | 2.48 | 3.68 | >10 | 4.59 | <100 |
|  |  |  | V2-Glycan |  |  |  | V3-Glycan |  | CD4bs | Interface | MPER | Autologous Plasma |
|  |  |  | PG9 | PG16 | PGT145 | CAP256.09 | AIIMS-P01 | PGT121 | VRC01 | PGT151 | 10-E8 |  |
| AIIMS_330 | 90-mo | 330.14.A12 | >10 | 5.23 | 1.73 | >10 | 0.33 | <0.02 | >10 | >10 | 0.21 | 912 |
|  |  | 330.14.A20 | 1.11 | >10 | >10 | >10 | 0.33 | 0.06 | >10 | >10 | 1.07 | 321 |
|  | 112-mo | 330.15.C36 | >10 | >10 | >10 | >10 | 0.779 | 0.551 | >10 | >10 | 6.73 | 148 |
|  |  | 330.15.D36 | >10 | >10 | >10 | >10 | 0.824 | 0.753 | >10 | >10 | 6.83 | <100 |
|  |  | 330.15.E36 | >10 | >10 | 2.41 | >10 | >10 | 1.43 | >10 | >10 | >10 | <100 |
|  | 117-mo | 330.16.A2 | >10 | >10 | >10 | >10 | 2.11 | 0.86 | >10 | >10 | 0.95 | 214 |
|  |  | 330.16.D4 | >10 | >10 | 4.37 | >10 | <0.02 | <0.02 | >10 | >10 | 1.51 | 151 |
|  |  | 330.16.E6 | 0.14 | 1.36 | <0.02 | <0.02 | 0.28 | 0.05 | 1.63 | 5.74 | 4.26 | 2459 |
|  |  | 330.16.H2 | 2.79 | >10 | 3.94 | >10 | 4.16 | 0.83 | >10 | >10 | 0.48 | 213 |

**Table 5:** Variable region characteristics of AIIMS_329 and AIIMS_330 in the timeframe of 90-months to 117-months. Variable loop length and PNGS of longitudinal SGA amplified amplicons of AIIMS_329 and AIIMS_330 were calculated using the variable region characteristics tool available at HIV Database (https://www.hiv.lanl.gov/). Kruskal-Wallis test was used to compare the difference in variable loop length and PNGS in the timeframe of 90- months to 117-months.

|  |  | Sequences (SGA Amplicons) | V1 |  | V2 |  | V3 |  | V4 |  | V5 |  |
| --- | --- | --- | --- | --- | --- | --- | --- | --- | --- | --- | --- | --- |
|  |  |  | Length | PNGS | Length | PNGS | Length | PNGS | Length | PNGS | Length | PNGS |
| AIIMS_329 | 90-months | 26 | 26 | 2 | 42 | 2 | 36 | 1 | 30 | 3 | 16 | 2 |
|  | 112-months | 24 | 28 | 5 | 56 | 2 | 36 | 1 | 37 | 6 | 12 | 2 |
|  | 117-months | 29 | 40 | 5 | 50 | 2 | 36 | 1 | 38 | 5 | 12 | 1 |
|  | p-value |  | **** | **** | **** |  |  |  | **** | **** |  |  |
| AIIMS_330 | 90-months | 24 | 24 | 4 | 44 | 3 | 37 | 1 | 37 | 5 | 17 | 2 |
|  | 112-months | 20 | 23 | 4 | 55 | 5 | 37 | 1 | 36 | 3 | 17 | 2 |
|  | 117-months | 21 | 34 | 4 | 59 | 5 | 37 | 1 | 36 | 5 | 16 | 2 |
|  | p-value |  | **** |  | **** | **** |  |  |  |  |  |  |

The sequences of SGA amplified envelope genes representing the circulating strains of both the twins at 90 -months were used to construct a phylogeny tree in order to identify the most recent common ancestor (MRCA) (Figure 3), which in turn was used as a reference sequence to generate highlighter plots (HIV Database). The highlighter plots for AIIMS_329 showed maximum divergence in V2C2, V3C3 and V4C4, with V4C4 envelope regions, with V4 loop lengthening, and acquiring the most PNGS (Potential N-linked glycosylation sites) in 112- and 117-months timepoint samples (Figure 4). In case of AIIMS_330, major divergence was observed in V2C2 and V4C4 regions of envelope glycoprotein in 112 and 117 time point samples (Figure 5). As V1V2 loops have been shown to be involved in virus escape from bnAbs, we aligned V1V2 loops of AIIMS_329 and AIIMS_330 to identify the key residues involved in resistance to known bnAbs as well as the autologous plasma. In case of AIIMS_329 virus, the key glycan residue N160 was mutated to Y, thereby leading to loss of N160 glycan. As the plasma V2-glycan directed antibodies in AIIMS_329 were solely dependent on N160, the N160Y mutation may be one of the potential escape mechanisms employed by AIIMS_329 viruses against contemporaneous plasma antibodies. In case of AIIMS_330, the key residues at N156, N160, K171 were preserved in the viruses resistant to autologous plasma antibodies, the resistant viruses however had longer V1 and V2 loops with increased PNGS (Figure 6). The viruses of AIIMS_329 probably developed resistance to autologous plasma antibodies by V1 and V4 loop lengthening (25,26) increased number of PNGS and loss of critical epitopes like N160 (Table 6 and Figure 6), features that have been associated with the development of resistance to bnAbs (27).

**Figure 3:**
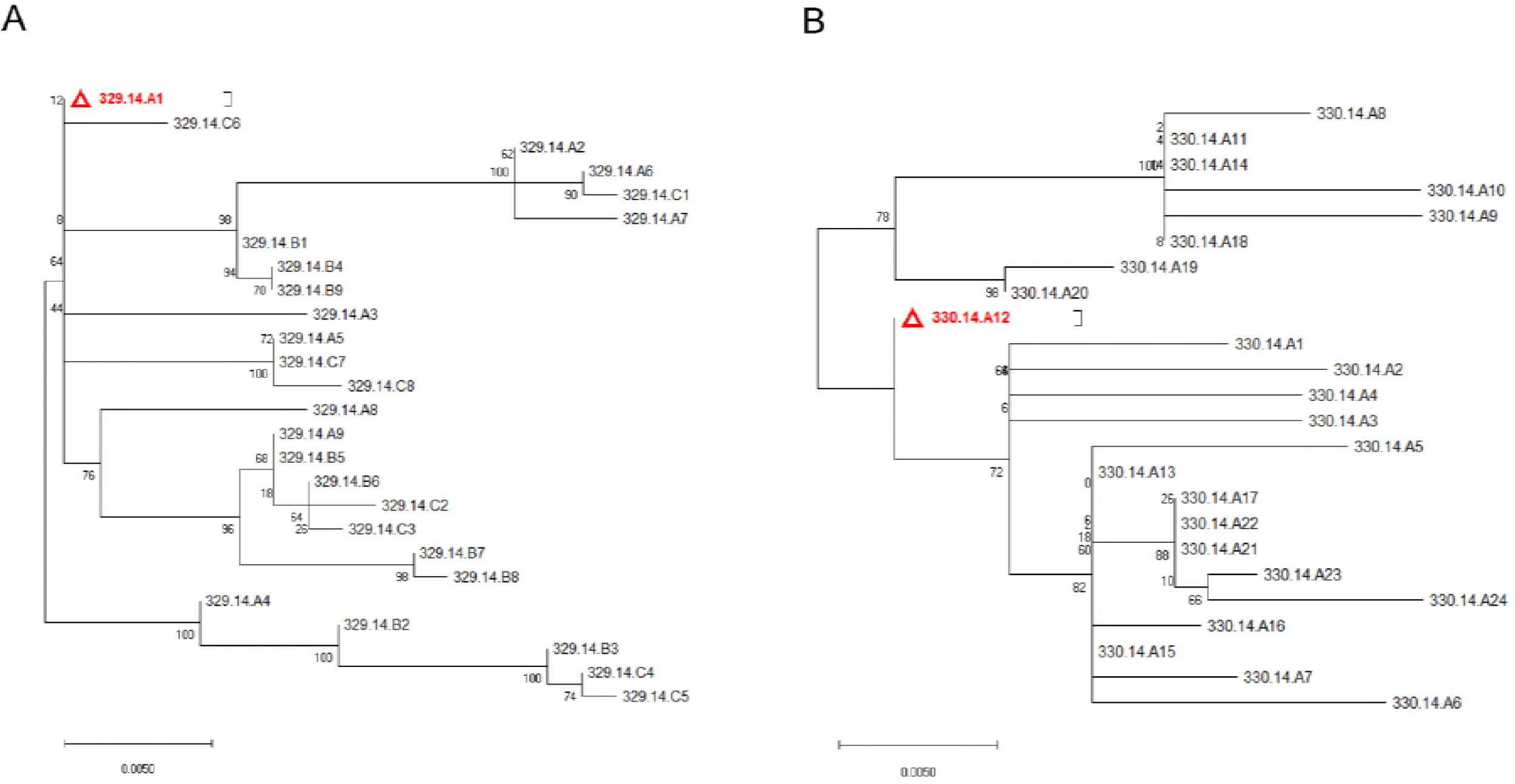
A neighbour-joining tree with 100 bootstrap replicates of all the SGA amplicons of AIIMS_329 (A) and AIIMS_330 (B) from 90-month timepoint was generated to identify the most recent common ancestor (MRCA) to be used as a reference sequence for generation of highlighter plots. The scale bar represents 0.0050 nucleotide substitution per site.

**Figure 4:**
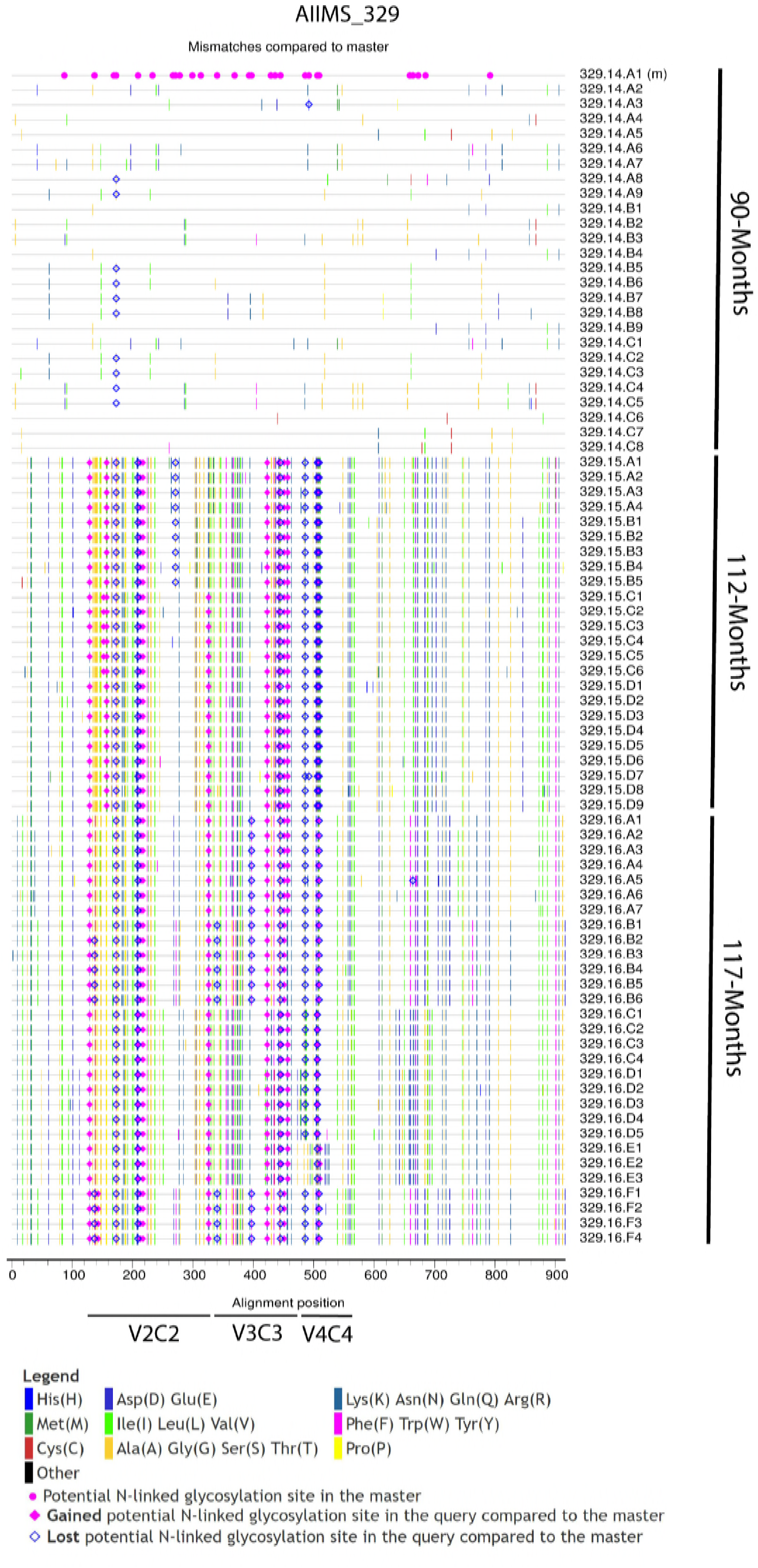
Highlighter plot of SGA amplicons of AIIMS_329. Using the most recent common ancestor from 90-month timepoint as a consensus, the highlighter plot shows amino acid mutations, gain and loss of potential N-linked glycosylation sites in longitudinal evolving viruses. The number on x-axis denoted the amino acid position, with V2C2, V3C3 and V4C4 sites marked and labelled with horizontal bars.

**Figure 5:**
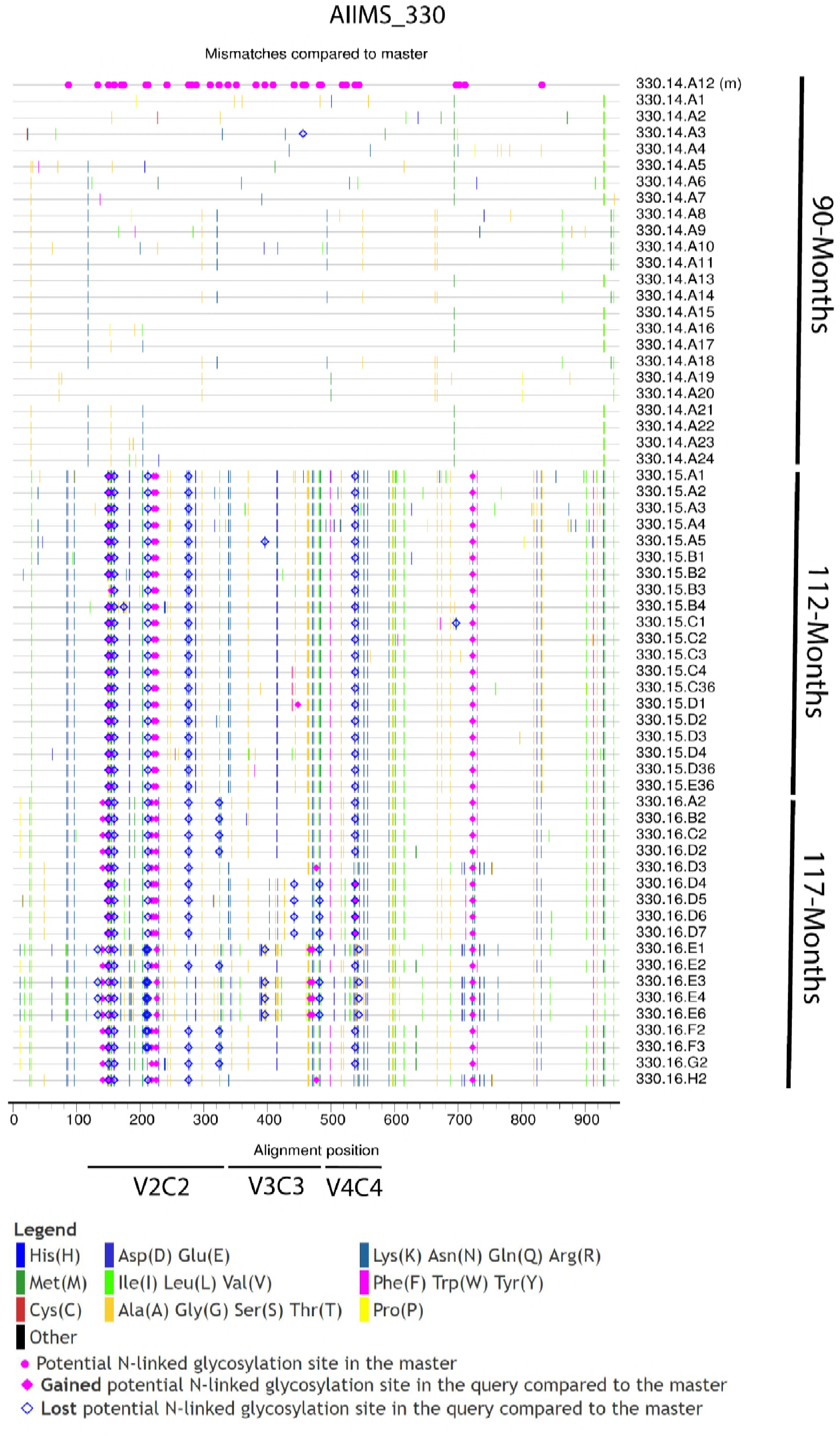
Highlighter plot of SGA amplicons of AIIMS_330. Using the most recent common ancestor from 90-month timepoint as a consensus, the highlighter plot shows amino acid mutations, gain and loss of potential N-linked glycosylation sites in longitudinal evolving viruses. The number on x-axis denoted the amino acid position, with V2C2, V3C3 and V4C4 sites marked and labelled with horizontal bars.

**Figure 6:**
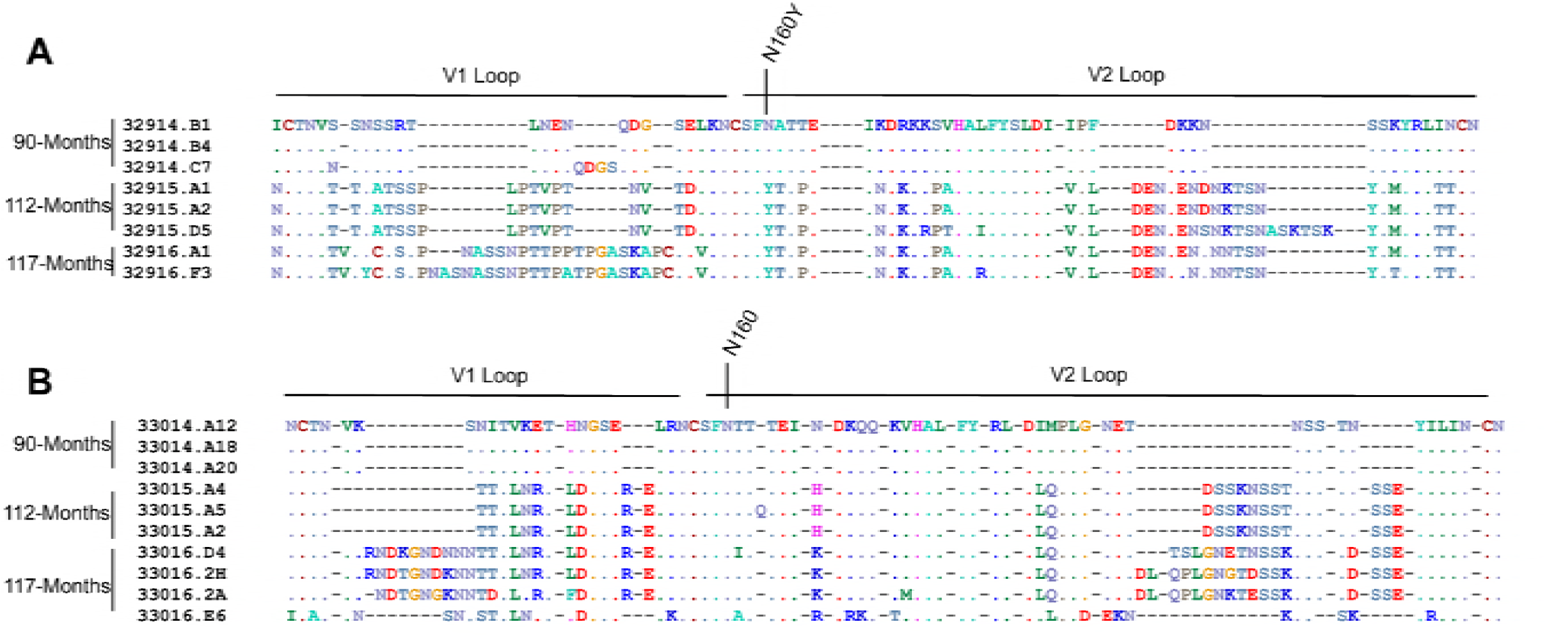
Alignment showing the sequence of V1V2 loops of AIIMS_329 (A) and AIIMS_330 (B) functional pseudoviruses, and distinct mechanism employed by them to escape autologous plasma neutralization. V1 and V2 loops are shown by the horizontal bar and N160, the site for V2-glycan, is labelled with the vertical bar.

## Discussion

Pediatric HIV-1 infection, with its biphasic mode of progression is distinct compared to the infection and pattern of disease progression in adults. Significantly high viral loads in context of normal CD4^+^ T cell counts, rapid disease progression, and generation of potent and polyclonal antibody response earlier in infection than adult counterparts are some of the differences between pediatric and adult HIV-1 infection. Recently, we and others observed the longitudinal evolution of plasma cross-neutralization antibody response with multiple epitope specificities in select antiretroviral naïve HIV-1 clade C (HIV-1C) chronically infected children (5,6). The clade C primary isolates that we established from some of these chronically infected children were relatively resistant to some of the tested second generation adult bnAbs (18). From this cohort of HIV-1 infected antiretroviral naive children that were longitudinally studied, we identified and characterised herein a pair of genetically identical twins, who were infected at the time of birth by vertical transmission. Both the twins were LTNPs at the time of recruitment. The presence of plasma bnAbs prompted us to further map the epitope specificities and viral diversity in this pair of identical twins with similar genetic and ethnic background, in order to understand the factors that contribute to the co-evolution of virus and plasma antibodies in antiretroviral naive chronically infected children of the same genetic makeup. We evaluated the neutralizing activity of plasma antibodies in this pair of HIV-1 infected twins over a period of 60 months, isolated and characterized functional pseudoviruses, and identified viral evolutionary mechanisms that plausibly may have been associated with development of plasma potency and breadth.

The plasma neutralizing activity of both twins was assessed over a period of 60 months, with the first sampling at 78-months post-infection (p.i.), using a global standard panel of 12 HIV-1 envelope pseudoviruses (21). A consistent increase in potency, median ID50 neutralization titre and GMT was observed for AIIMS_330 plasma, that peaked at 112-months post infection (p.i), and then reached a plateau throughout the duration of the study. AIIMS_329, on the other hand, showed highest GMT, median ID50 neutralization and potency at 90-months p.i, which then declined at 112-months p.i, beyond which the titres stabilized. In both HIV-1 infected adults and children, a continuous increase in breadth and potency of plasma antibodies is well established (6,28,29), however a reduction in breadth and potency of the AIIMS329 plasma neutralizing antibodies at 112-months p.i, while maintaining a stable CD4^+^ T cell count and controlled viremia, was intriguing. The lower viral load in AIIMS_329 as compared to AIIMS330 and a strain specific enrichment of plasma antibodies at 112-months p.i may have contributed to the reduction in neutralization breadth in AIIMS_329. A longitudinal analysis of the plasma neutralization breadth and potency of both twins upto 112-months p.i against a multiclade panel of 45 viruses, including 10 viruses of Indian origin, further confirmed that the AIIMS_329 plasma antibodies were progressively less effective in cross neutralizing activity while retaining their ability to neutralize viruses of Indian origin. The AIIMS_330 plasma showed elite neutralizing activity from 78-months p.i, which may have developed due to the persistent antigenic stimulus by the high viremia seen in this twin as compared to AIIMS_329. Long exposure to high viral load with declining CD4^+^ T cell counts has been earlier shown to be associated with development of plasma breadth and potency over time (28).

The epitope specificities of plasma bnAbs for AIIMS_329 and AIIMS_330 were assessed by performing neutralization assays using N160A V2-glycan mutants, N332A V3-glycan (mutants, CD4bs probe RSC3 and its mutant RSC3Δ371I/P363N, as well as MPER peptides MPER-B and –C. AIIMS-330 plasma showed dependence on N160A glycan at 78-month p.i (baseline sample) which increased in potency over time (From 4.1 to 8.9), whereas AIIMS-329 plasma showed N160A glycan dependence 102-months p.i, two years later than AIIMS-330. No reduction was observed with 16055 virus and its mutants N156K, K169A and K171A, indicating that the V2-glycan dependence was mostly on the N160 epitope in V2 apex. Both the twins showed V3-glycan dependence at 112-months p.i. At 112-month, AIIMS-330 plasma also developed CD4bs dependence whereas AIIMS-329 failed to generate CD4bs dependence throughout the duration of the study. Neither twin generated MPER dependence. Both the twins developed primary neutralizing response against V2 glycans to begin with, which is in accordance with several studies showing V2 glycan bnAbs to be most common and potent (29,30). Many studies have shown that children mount broader and more potent response against HIV-1 earlier than adults with one report showing the presence of broad neutralizing activity in one infant one-year p.i (4,7). Polyclonal bnAb responses in HIV-1 infection have been well reported in instances of both adult and pediatric infection (31–33), with chronic infection and higher antigenic stimulation, often identified as factors responsible for neutralization dependence on multiple specificities. Our observation of dual epitope specificity in the AIIMS-329 plasma and antibodies against 3 distinct epitopes in AIIMS-330 plasma that developed by 112-months of age is in consonance with the findings of a recent study wherein children infected with HIV-1 for a duration of ten years on average were found to have multiple HIV-1 nAb specificities (5). Based on the multiple epitope specificities observed in AIIMS-330 plasma 112-months p.i, and the highest GMT titres and median ID50 values obtained from the contemporaneous plasma mapping analysis using the 45-virus panel, we inferred that the spike observed in AIIMS-330 plasma BCN activity at the fifth follow-up was probably due to the development of V3-glycan and CD4bs dependence at the same timepoint. Despite the presence of plasma neutralization antibodies with multiple epitope specificities, and having similar antigenic exposure, genetic and ethnic background, both twins continued to have viremia and developed different plasma neutralization profile and dependency with time.

Both the virus and antibody responses in the host are known to co-evolve, and a continuous interplay between HIV-1 envelope specific antibody response and viral escape is one of the key features of bnAb development (12,34–39). In order to trace the viral evolution in these twins that occurred along with the increase in the plasma antibody neutralizing activity with time, we assessed the circulating viral strains in both the twins around the timepoint (90-months to 117-months) when they developed multiple specificities to delineate the viral properties that may be associated with the altered plasma neutralization potency. To negate the cloning bias often associated with conventional bulk PCR amplification approach, functional pseudoviruses representing circulating HIV-1 strains were generated via single genome amplification of envelope genes, from HIV-1 RNA isolated from plasma samples of the twins and their susceptibility to neutralization by contemporaneous plasma as well as bnAbs was evaluated. To further strengthen that the functional pseudoviruses generated represented the currently circulating viral strains, we sequenced more than 20 SGA PCR amplicons from each time point studied with a 90% confidence (40). In AIIMS_330 plasma, we observed diverse circulating viral strains; viruses that showed neutralization susceptibility to autologous plasma antibodies and bnAbs as well as resistant viruses, throughout the timeframe of 90-months to 117-months p.i. In AIIMS_329, the viruses generated at 90-month p.i were susceptible to majority of the bnAbs as well as contemporaneous autologous plasma antibodies, with the viruses generated from 112-months and 117-months p.i. resistant to bnAbs and contemporaneous autologous plasma. We observed two distinct evolutionary pathways of viral escape in response to increasing plasma potency and breadth in the twins. The viruses of AIIMS_329 developed resistance to autologous plasma by previously established mechanisms (25–27,41,42) of the V1, V2 and V4 loop lengthening, increased number of PNGS and loss of critical epitopes like N160. The AIIMS_330 viruses that were resistant to contemporaneous autologous plasma had longer V2 loop but maintained all epitopes. In AIIMS_329, all the viruses sampled at 112-months and 117-months p.i. were resistant while AIIMS_330 viruses formed a pool of viruses with some sensitive and others resistant to the contemporaneous autologous plasma antibodies. In the donor from which the CAP256-VRC26 lineage of antibodies were isolated (12,35,43), the CAP256-VRC26 lineage resistant primary infecting virus (CAP256-PI), present along with the lineage sensitive superinfecting virus (CAP256-SU), drove the evolution of CAP256-VRC26 lineage towards antibodies with increased breadth and potency. On similar lines, we observed that the plasma neutralizing antibodies of AIIMS_330 donor, with a mixture of sensitive and resistant viruses, had high potency, GMT, and multiple antibody specificities, suggesting a correlation between the virus pool with varied susceptibility profile and the development of plasma breadth and potency. Further we observed that the viruses sampled from AIIMS_329 at the time point when plasma potency decreased (112-months) were less susceptible to neutralization by autologous plasma antibodies as well as major bnAbs.

To summarize, in this study, we describe for the first time, the diverse evolution of plasma neutralizing antibodies in a pair of genetically identical twins over a period of 60-months, despite the same time point of infection via a similar virus and similar CD4^+^ T cell counts with an added advantage of matched timepoint assessments between two infected individuals belonging to one transmission pair. Our findings are in corroboration with previously published reports suggesting that loop lengthening with increased glycan density contributes to the development of viral resistance to V2-apex bnAbs. In addition, we observed potential viral evolution in AIIMS_330 donor, harbouring a viral pool constituting autologous plasma resistant and sensitive viruses that may have contributed to the development and maintenance of elite neutralizing activity. The data emerging from the study of evolving viruses and plasma antibodies in this pair of HIV-1C chronically infected identical twins is supportive of envelope-based lineage vaccine approaches that are currently being evaluated. This strategy is based on use of a mixture of immunogens with different antibody imprinting ability rather than one immunogen with susceptibility to all bnAbs, that could provide better antigenic exposure, higher antigenic variability for enhanced somatic hypermutation, and may lead to induction of antibodies with increased potency and breadth and our data is supportive of this strategy.

## Materials and methods

### Study Population

AIIMS_329 and AIIMS_330 were recruited from Outdoor Patient Department of Department of Pediatrics, AIIMS for this study at the age of nine years and were followed for a total period of 60 months. Blood was drawn in 5-ml EDTA vials, plasma was aliquoted for plasma neutralization assays and viral RNA isolation, and viral loads. The study was approved by institute ethics committee of All India Institute of Medical Sciences (IEC/NP-295/2011 & RP-15/2011).

### Plasmids, viruses, monoclonal antibodies, peptides, and cells

Plasmids encoding HIV-1 env genes representing different clades, monoclonal antibodies and TZM-bl cells were procured from NIH AIDS Reagent Program. Plasmid encoding RSC3 wild type probe and its mutant RSC3Δ371I/P363N were kindly provided by John Mascola, National Institute of Allergy and Infectious Diseases, USA. Plasmids for CAP256 and its N332A mutant was kindly provided by Lynn Morris, National Institute of Communicable Diseases, SA. MPER-B and -C peptides were commercially synthesized from Sigma Genosys, USA (23,24). HEK293T cells were purchased from American Type Culture Collection (ATCC).

### Cloning of autologous HIV-1 envelope genes and production of replication incompetent pseudoviruses

Autologous replication incompetent envelope pseudoviruses were generated from AIIMS_329 and AIIMS_330 as described previously (44). Briefly, viral RNA was isolated from 140 µl of plasma using QIAamp Viral RNA Mini Kit, reverse transcribed, using gene specific primer OFM19 and Superscript III reverse transcriptase, into cDNA which was used in two-round nested PCR for amplification of envelope gene using High Fidelity Phusion DNA Polymerase (New England Biolabs). The envelope amplicons were purified, and ligated into pcDNA3.1D/V5-His-TOPO vector (Invitrogen). Pseudoviruses were prepared by cotransfecting 1.25 µg of HIV-1 envelope containing plasmid with 2.5 µg of an envelope deficient HIV-1 backbone (PSG3Δenv) vector at a molar ratio of 1:2 using PEI-MAX as transfection reagent in HEK293T cells seeded in a 6-well culture plates. Culture supernatants containing pseudoviruses were harvested 48 hours post-transfection, filtered through 0.4µ filter, aliquoted and stored at -80°C until further use. TCID50 was determined by infecting TZM-bl cells with serially diluted pseudoviruses in presence of DEAE-Dextran, and lysing the cells 48 hours post-infection. Infectivity titres were determined by measuring luminescence activity in presence of Bright Glow reagent (Promega).

### HIV-1 envelope sequences and phylogenetic analysis

HIV-1 envelope genes were PCR amplified from plasma viral RNA by single genome amplification (40) and directly sequenced commercially. Individual sequence fragments of SGA amplified amplicons were assembled using Sequencher 5.4 (Gene Code Corporation). Nucleotide sequences were aligned in MEGA X (45) and phylogeny trees for identifying the most recent common ancestor were constructed by the neighbour-joining method with 100 bootstrap replicates (44).

### Neutralization assay

Neutralization assays were carried out using TZM-bl cells, a genetically engineered HeLa cell line that constitutively expresses CD4, CCR5 and CXCR4, and contains luciferase and β-galactosidase gene under HIV-1 tat promoter, as described before (46). Briefly, envelope pseudoviruses were incubated in presence of serially diluted bnAbs, or heat inactivated plasmas for one hour. After incubation, freshly Trypsinized TZM-bl cells were added, with 25

µg/ml DEAE-Dextran. The plates were incubated for 48 h, cells were lysed in presence of Bright Glow reagent, and luminescence was measured. Using the luminescence of serially diluted bnAbs or plasma, a non-linear regression curve was generated and titres were calculated as the bnAb concentration, or reciprocal dilution of serum that showed 50% reduction in luminescence compared to untreated virus control. For V2- and V3-glycan dependence, pseudoviruses with key mutation in V2- and V3-glycan were used with their wild type counterparts, and were incubated with plasma for one hour at 37°C followed by addition of TZM-bl cells, and readout 48-hour post infection.

### Binding ELISAs

MPER, RSC3 and RSC3Δ371I/P363N ELISAs were performed as described previously (16). Briefly, 96 well ELISA plates (Corning, USA) were coated with 2 µg/ml of RSC3, and RSC3Δ371I/P363N proteins and MPER-B and MPER-C peptides overnight at 4°C. Coated plates were washed with PBS containing 0.05% Tween 20. Plates were blocked with 5% skimmed milk in blocking buffer. A 50-fold dilution of plasmas, was added, titrated, and incubated at 37°C for I hour. Unbound plasma antibodies were washed with wash buffer and plates were incubated with peroxidase conjugated goat anti-human IgG at a dilution of 1:1000. Following secondary antibody incubation, the wells were washed, and TMB substrate was added. After color development, reaction was stopped with 0.2 M H_2_SO_4_ and absorbance was measured at 450 nm.

### Nucleotide sequence accession numbers

The SGA amplified HIV-1 envelope sequences are available at GenBank with accession number MK076582-MK076724.

### Statistical analysis

The Mann Whitney U test was used for comparison of two unpaired parameters, Wilcoxon matched-pairs signed rank test was used for comparison of two paired parameters. Kruskal-Wallis test was used for comparison of three parameters. All statistical analyses were performed on GraphPad Prism 6. A p-value of <0.05 was considered significant.

## Acknowledgement

This work was funded by Department of Biotechnology, India (BT/PR5066/MED/1582/2012). The Junior Research Fellowship to N.M was supported by University Grants Commission (UGC), India. The Senior Research Fellowship to M.A.M was supported by Indian Council of Medical Research (ICMR), India. We thank AIIMS_329 and AIIMS_330 for participating in this study. We are thankful to NIH AIDS Reagent program for providing HIV-1 envelope pseudovirus plasmids, bnAbs and their plasmids, TZM-bl cells, and Neutralizing Antibody Consortium (NAC), IAVI, USA for providing bnAbs. We are thankful to Dr. John Mascola for providing RSC3 probe and its mutant, Dr. Lynn Morris for providing V2- and V3-glycan mutant plasmids.

## Author contributions

N.M, M.A.M, and K.L designed the study, analyzed data, wrote and edited the manuscript. N.M and M.A.M planned, performed the experiments, and analyzed data. S.S and D.K contributed to SGA amplification, and neutralization assays. S.K contributed to binding ELISAs, and neutralization assays. H.C contributed to binding ELISAs. U.K performed HLA Haplotyping, and edited the manuscript. R.S, R.L, S.K.K, B.K.D contributed and coordinated AIIMS_329 and AIIMS_330 samples.

